# Harnessing cellular perception apparatus for smart metabolic reprogramming

**DOI:** 10.1101/2022.04.03.486851

**Authors:** Chunlin Tan, Fei Tao, Ping Xu

## Abstract

Metabolic reprogramming (MRP) is a fundamental approach in synthetic biology that involves redirecting metabolic flux and remodeling metabolic networks. However, only few approaches have been made in effective metabolic operations, especially at global level of metabolic networks. Naturally existing cellular perception apparatuses (CPAs), such as histidine kinases (HKs), are considered to sit on sensitive nodes of the metabolic network, which can trigger natural MRP upon perceiving environmental fluctuations. We develop a plateform for global MRP by natural environmental stimulation based on the combinational interference of CPAs. The plateform consists of a CRISPRi-mediated dual-gene combinational knockdown (CDCK) strategy and survivorship-based metabolic interaction analysis (SMIA). A total of 35 histidine kinase (HK) genes and 24 glycine metabolism genes were selected as targets to determine effectiveness of our approach for fast-growing chassis *Vibrio* FA2. Combined knockdown of several genes of HKs and glycine metabolism increased the glycine production. Other other hand, effects of CDCK on bacterial antibiotic resistance were assessed by targeting HKs. Many HKs were identified to be associated with antibiotic resistance in *Vibrio* FA2, of which combinational knockdown of two HK genes *sasA_8* and *04288* reduced the ampicillin resistance. This MRP strategy is powerful and cost-effective, and can be considered as a smart strategy capable of operating a broad range of metabolic networks in microorganisms.

## Main

Synthetic biology, an interdisciplinary area of engineering and science in the 21st century, is very helpful in exploring the fundamental laws of life activities and breakthroughs in biotechnology innovation ^1^. Accordingly, novel biomolecular components, networks, and pathways can be designed and built to reprogram organisms ^2^, a process also called metabolic reprogramming (MRP) through its definition. MRP is a fundamental approach in synthetic biology that involves the redirection of metabolic flux and remodeling of the metabolic networks ^3^. For example, the yeast cell growth on xylose was substantially enhanced by performing a global rewiring of the cellular metabolic network ^4^. Wang et al. discovered a switch in metabolic status that can reshape the cellular metabolic network, resulting in high-level acetoin production ^5^. MRP has also been recognized to be involved in cell differentiation, cancer progression, and metastasis ^6, 7^. Antibiotic resistance has been repeatedly reported to result from spontaneous MRP ^8-10^. Therefore, MRP is a crucial handle for controlling and understanding cells, both in biophysiology and biomanufacturing.

Unfortunately, it is difficult to artificially achieve an effective MRP, especially at global level of the metabolic network. Theoretically, the primary obstacle relies on the high complexity of the chaotic metabolic network, which consists of thousands of metabolites, enzymes, genes, and their interactions ^11^. As this complexity results in poor predictability of metabolic behavior, it is difficult to infer the final cellular response to a given metabolic perturbation ^12^. Practically, the lack of an effective approach for metabolic interference severely limits the space of trials in MRP studies, depriving the advanced opportunity for the MRP research ^13^. Currently, it is still time-consuming and labor-intensive to obtain the desired production performance in the biological industry ^14^. For example, after more than ten years of hard work and more than 50 million US dollars, Paddon et al. industrialized the biosynthesis process of artemisinin and achieved large-scale mass production of artemisinic acid ^15^. Therefore, there is an urgent need to develop powerful tools for addressing these challenges.

Various environmental cues, such as light, temperature, chemicals, pH, and pressure, are well known to naturally stimulate microbial cells to switch on their cell MRP in response to these changes ^16^. This natural MRP is achieved by sensing environmental signals through cellular perception apparatuses (CPAs), which are widely distributed on the cell surface (Fig. 1). As typical CPAs, histidine kinases (HKs) perceive and transduce many environmental signals to trigger cellular MRP corresponding to many crucial physiological processes such as cell development, antibiotic resistance, and chemotaxis ^16^. Moreover, transcription factors (TFs) with sensor domains can switch cellular metabolic status by interacting with small (effector) molecules, ions, physical parameters (e.g. temperature or pH), amino acids, succinate, and secondary metabolites ^17-20^. Commonly, natural MRP encoded by CPAs is substantial and effective owing to their extreme importance for the survival of cells in fluctuating environments ^21^. Therefore, simulating the natural MRP caused by environmental vibrations via CPA manipulation would be an effective strategy for reprogramming microbial metabolic networks.

**Fig. 1.**
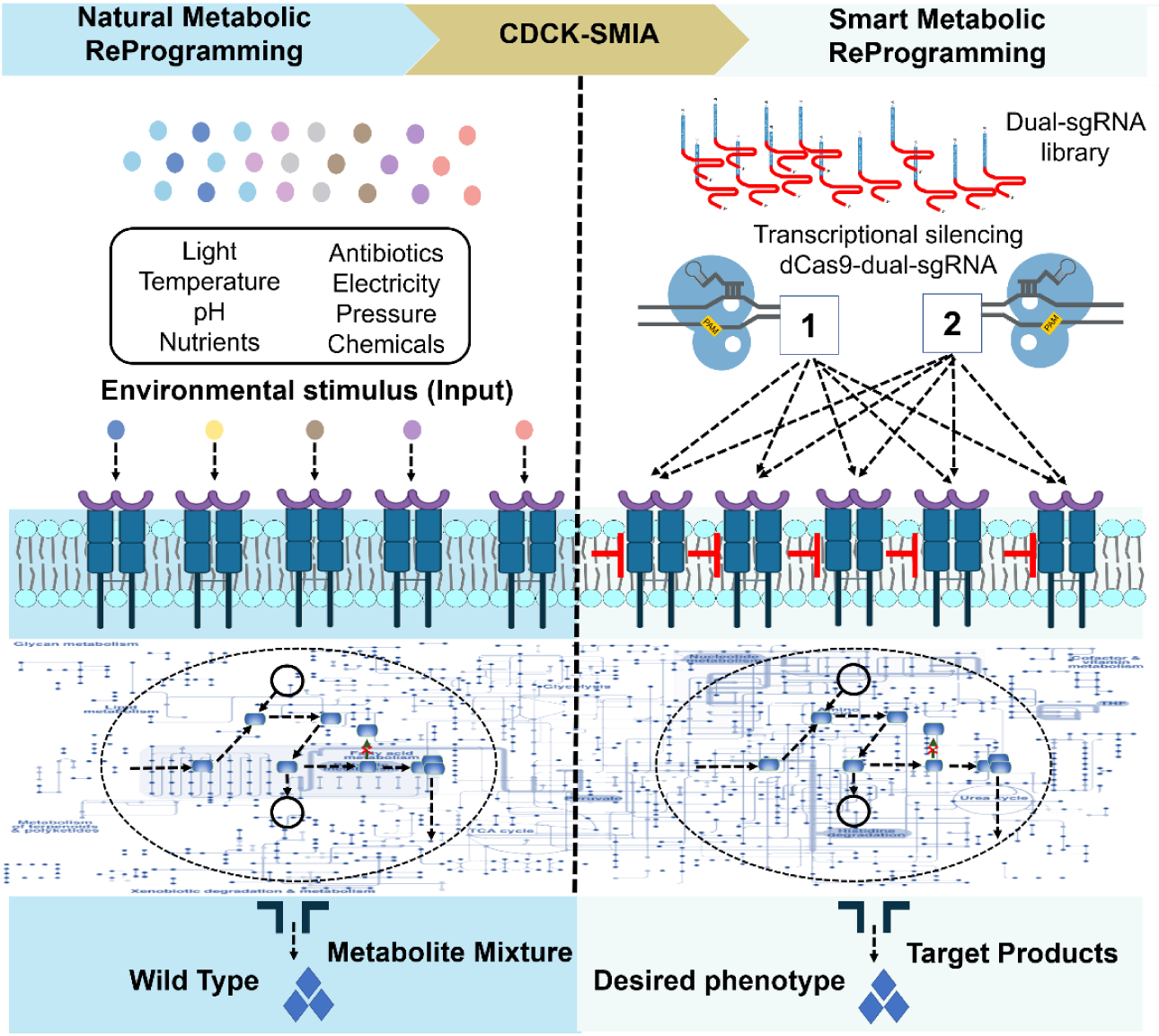
Schematic diagram of natural metabolic reprogramming and smart metabolic reprogramming. The different environmental cues, such as light, temperature, chemicals, pH, and/or pressure, naturally stimulate microbial cells to switch on their cell metabolic reprogramming (MRP) in response to these changes. The naturally existing cellular perception apparatuses, such as histidine kinases (HKs), are considered sitting on sensitive nodes of the metabolic network, which can trigger natural MRP upon perceiving environmental fluctuations. This study develops a platform for global MRP by the natural environmental stimulation based on combinational interference of cellular perception apparatuses (CPAs). The well-developed method is exceptionally effective in changing metabolic status, which can be considered smart MRP in terms of cost, convenience, and throughput. CDCK-SMIA: the CRISPRi-mediated dual-gene combinational knockdown (CDCK) strategy coupled with survivorship-based metabolic interaction analysis (SMIA).

In this study, we developed a platform that can harness CPAs for MRP from a global metabolic perspective in a *Vibrio* strain FA2 (Fig. 2a). This platform consists of a CRISPRi-mediated dual-gene combinational knockdown (CDCK) strategy and survivorship-based metabolic interaction analysis (SMIA; Fig. 2b, 2c, and 2d). Using this platform, we investigated the effects of MRP targeting HKs and glycine pathway genes on glycine production by *Vibrio* FA2. In addition, we determined the effects of CDCK on bacterial antibiotic resistance by targeting HKs and selected four genes for verification with wet experiments. The results of wet experiments were consistent with those of the bioinformatics analysis. Overall, our platform is powerful and cost-effective and can be considered as a smart MRP strategy that can be applied to operate a broad range of microbial metabolic networks.

**Fig. 2.**
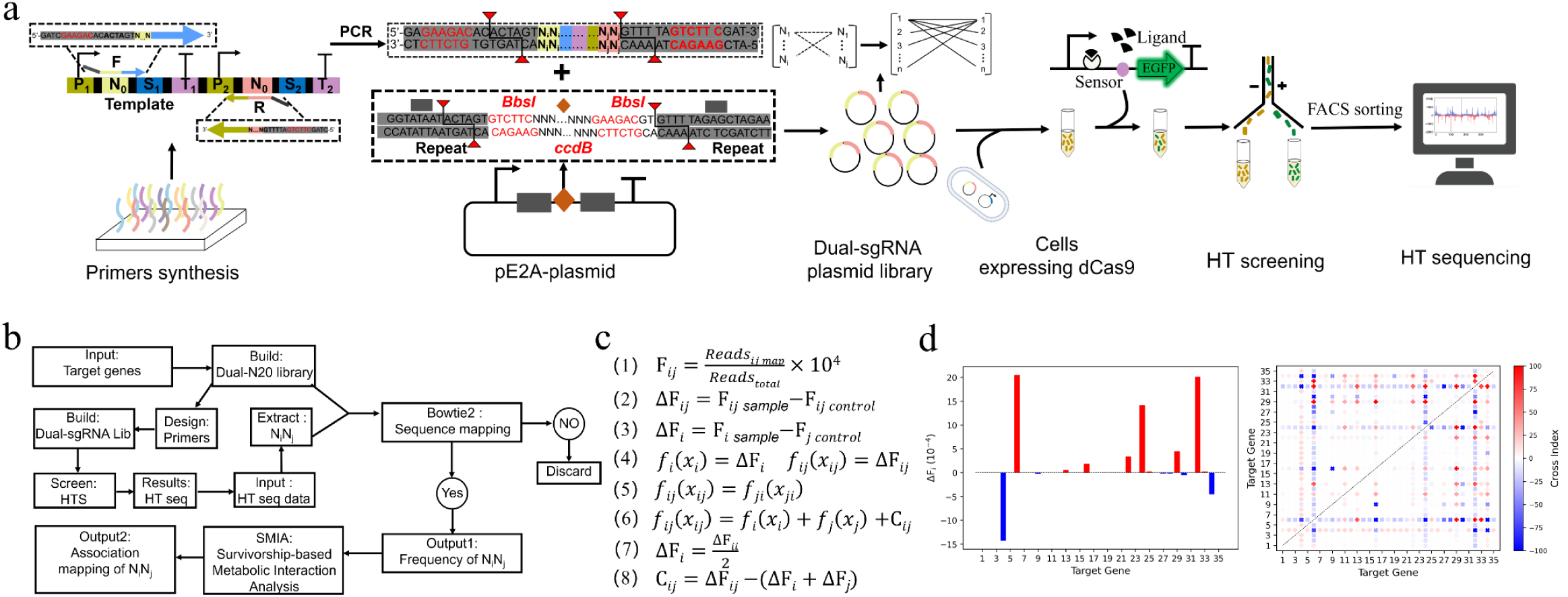
Overview of this study. **a** Schematic representation of the CRISPRi-mediated dual-gene combinational knockdown (CDCK) strategy coupled with the biosensor-mediated high-throughput (HT) screening processes. A dual-sgRNA library targeting genes of interest is constructed by the polymerase chain reaction (PCR) and Golden Gate Assembly. The dual-sgRNA library gene segments are amplified from the template and cloned into the pE1a-tool plasmid by the Golden Gate Assembly, transformed into *E. coli*, resulting in dual-sgRNA library plasmid-producing strains. The dual-sgRNA plasmid is further transformed into *Vibrio* FA2 expressing dCas9 protein and glycine-on riboswitch mediated eGFP protein, resulting in cell libraries. The cell libraries are grown under isopropyl-β-d-thiogalactoside (IPTG) and arabinose to induce the eGFP protein and dCas9 protein, respectively. The induced cell libraries are sent into fluorescence-activated cell sorting (FACS) and higher fluorescence signal cells were screened out compared to the control. The plasmids of sorted cells are extracted as the template. Then, the dual-sgRNA segments are amplified by PCR from the template and are sent to next-generation sequencing (NGS). **b** Flow diagram of the CDCK strategy design and survivorship-based metabolic interaction analysis (SMIA) for the smart metabolic reprogramming (MRP) platform. In this study, we designed a website providing a one-stop service for the smart MRP research. It allows a user to automatically design related experiments. After the wet experiments, one can use the services provided to conduct data analysis by simply uploading the raw data. **c** Calculation equations of the SMIA platform. Defined i, j and C as the gene number, and cross index. In theory, position effects of the i and j gene are the same, as shown in equation (5). Equation (7): C_ij_ = 0, i = j. Equation (8): C_ij_ ≠ 0, i≠j. All of the data was experimentally obtained except for equation (7), which was inferred from the data. **d** Representative results of the SMIA platform. (Left) Contribution of the single knockdown gene. (Right) Contribution of the combinational knockdown gene. (If C_*ij*_ > 0, it indicates that i and j genes have a positive interaction, showing “+”. If C_*ij*_=0, then the i and j genes are not interacting with each other, showing in white. If C_*ij*_ < 0, then the i and j genes interact negatively, displaying as “×”.) The ΔF_i_ indicates the contribution of single gene knockdown to the survival rate of whole-cell libraries.

## Results

### Construction and testing of CRISPRi-mediated multiple gene knockdown

To test functionality of the CRISPRi system in *Vibrio* FA2, we created a reporter *Vibrio* FA2 expressing mRFP (monomeric red fluorescent protein ^22^), a catalytically dead Cas9 (dCas9 ^23^), and sgRNAs complementary to different regions of the mRFP coding sequence ^23^, either the template (T) or nontemplate (NT) DNA strand (Fig. 3a). sgRNAs targeting the non-template DNA strand could effectively repress mRFP gene expression (> 10-fold repression), which is was consistent with the microscopic image results. We proceeded to determine whether the CRISPRi system could function well in controlling multiple genes independently without crosstalk in *Vibrio* FA2. Briefly, a dual-color fluorescence system based on mRFP and eGFP (enhanced red fluorescent protein ^22^) was devised, and then two sgRNAs complementary to the NT1 of the mRFP and eGFP coding sequences were designed (Fig. 3a). The expression of each sgRNA was only found to silence the cognate gene and had no effects on the other genes. However, co-expression of the two sgRNAs knocked down both genes of *mrfp* and *egfp* (Fig. 3b and 3c).

**Fig. 3.**
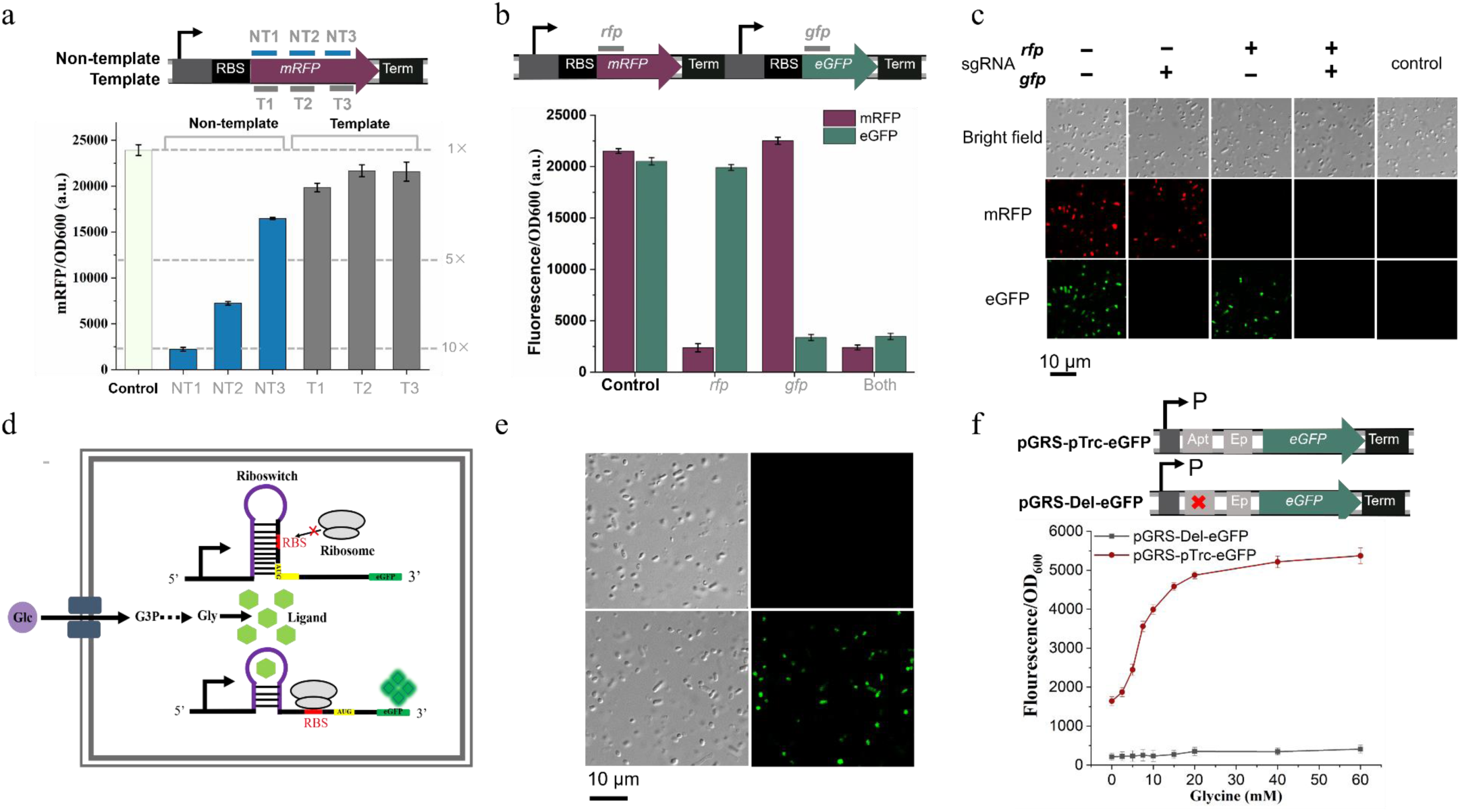
CRISPRi-based multiple gene knockdown and glycine-on riboswitch function in the fast-growing *Vibrio* FA2 chassis. **a** CRISPRi blocks transcription elongation in a strand-specific manner. A synthetic fluorescence-based reporter system containing the mRFP-coding gene and eGFP-coding gene is used to test the CRISPRi system. Six sgRNAs that bind to the nontemplate (NT) DNA strand and template (T) DNA strand is expressed in the pE1a plasmid, with their effects targeting mRFP and eGFP measured by in vivo fluorescence assay. Only the sgRNAs that bind to the nontemplate showed silencing. The control shows fluorescence of the cells with dCas9 protein but without sgRNA. **b** Multiple sgRNAs can independently silence mRFP and eGFP fluorescence protein reporters in the same cell. Each sgRNA specifically silences its cognate gene but not the other gene. Both of the mRFP and eGFP-coding genes are repressed when both the sgRNAs are present. **c** Microscopic images for using two sgRNAs to control the fluorescence protein expression. (Top) Bright-field images of the *Vibrio* FA2 cells; (middle) RFP channel; (bottom) GFP channel. Co-expression of one sgRNA and dCas9 protein only silences the cognate fluorescent protein, but not the other. The knockdown effects are obvious as almost no fluorescence is observed from cells when targeting certain fluorescence protein genes. Scale bar, 10 μm. Control shows cells without fluorescence reporters. **d** Schematic diagram of the glycine-on riboswitch principle. In the glycine biosensor, eGFP can serve as a report signal so that the concentration of glycine is coupled to the fluorescence value. **e** Fluorescence microscopy images of *Vibrio* FA2 carrying the glycine-on riboswitch fed with (lower row) or without (upper row) the addition of glycine. **f** Fluorescence values of two different constructs. Relative eGFP expressions were measured in the absence of different glycine concentrations. pTrc: pTrc promoter; Apt: aptamer; Ep: expression platform. All data are averages of three independent experiments.

Based on these results, the CRISPRi system works well and sgRNA-guided targeting is specific and not affected by other sgRNAs in *Vibrio* FA2. Hence, CRISPRi-based multiple gene knockdown tools function well in fast-growing *Vibrio* FA2 chassis.

### Design and characterization of glycine-on riboswitch

Riboswitch-mediated biosensors are often powerful tools for timely and precise monitoring of compounds ^24^. A glycine-on riboswitch from the upstream nucleotide of the VC1422 gene of *Vibrio cholerae* ^25^ was used to develop a high-throughput screening (HTS) approach. A schematic of the principle of the biosensor is shown in Fig. 3d. The biosensor consists of two sections: i) a glycine-on riboswitch (containing two tandem aptamers and an expression platform) that could change its secondary structure and activate expression of the downstream gene in response to the increased amount of glycine, and ii) an eGFP that acts as a reporter gene for a selection marker of HTS. To test its feasibility in *Vibiro* FA2, a glycine-on riboswitch with different promoters was used to construct plasmids pGRS-eGFP, pGRS-J23100-eGFP, and pGRS-Native-eGFP, and the control plasmids (in which glycine aptamers were deleted) pGRS-Del-eGFP, pGRS-J23100-Del-eGFP, and pGRS-Native-Del-eGFP.

To determine whether the glycine-on riboswitch could respond to increased glycine concentrations, constructs and their corresponding control plasmids were transformed into *Vibrio* FA2. A previous study showed that a high glycine concentration might inhibit cell growth owing to the potential toxicity of excess glycine ^25^. To avoid inhibition, we determined the growth of *Vibrio* FA2 in a chemically defined optimized M9 medium ^26^ supplemented with various concentrations (10−100 mM) of glycine. Cell growth was found to be significantly inhibited in the optimized M9 medium supplemented with 70−100 mM glycine. Therefore, we tested the functionality of the glycine-riboswitch in *Vibrio* FA2 using an optimized M9 medium supplemented with various concentrations (0−60mM) of glycine. The dose-response of the glycine-on riboswitch-based eGFP sensors is shown in Fig. 3c and 3d. Our findings suggest that only pGRS-pTrc-eGFP can upregulate the eGFP gene expression in response to increased glycine concentration. In contrast, activation was not observed in the pGRS-pTrc-Del-eGFP or other constructs. The average eGFP fluorescence value sharply increased (> 3-fold) when the glycine concentration was increased from 0 to 60 mM (Fig. 3f). The glycine-riboswitch even slightly upregulated the downstream eGFP gene in the presence of 2.5 mM glycine. Owing to its sensitivity and large dynamic range, the glycine-on riboswitch was selected for the subsequent study. Overall, the glycine-on riboswitch functioned well as a switch for HTS in *Vibrio* FA2.

### Establishment of the CDCK-SMIA platform

To implement an MRP platform, together with the above results, we synthesized primers and constructed a dual-sgRNA library containing approximately 40,000 transformants in *E. coli* through PCR amplification and golden-gate assembly. Figure 2a shows a schematic diagram of our CDCK-SMIA plateform (see the details in the Methods section). The sgRNA library plasmid was then transferred into *Vibrio* FA2 containing dCas9 and pGRS-pTrc-eGFP expressed cassette by electroporation, resulting in approximately 40,000 transformants. The library was named in *E. coli* L1 and dCas9-containing *Vibrio* FA2 L2. In total, dual-sgRNA libraries targeting HKs (HK-L1 and HK-L2) and glycine pathways (defined as G-L1 and G-L2), respectively, were constructed. The L2 library was then cultivated in the LB3 medium supplemented with arabinose and IPTG to induce the CRISPRi system. Based on their fluorescence values, different cells from the CRISPRi-mediated dual-sgRNA library were screened using fluorescence-activated cell sorting (FACS) technology (sorted cells labeled S). The plasmids of these cells were extracted and the and dual-sgRNA cassette fragments were amplified using PCR since it contains the diversity of the cells. The PCR products were sent for NGS ^27^. In addition, the frequencies of dual-sgRNA cassettes in surviving cells were calculated and a survivorship-based metabolic interaction analysis (SMIA) was established. Therefore, the CDCK-SMIA platform was used for further experiments.

As shown in Fig. 2c, we defined i/or j, ΔF_i,_ and C_ij_ as the gene number, frequency of single-gene contribution, and frequency of dual-gene contribution (defined as cross index), respectively. F is the frequency value obtained experimentally or inferred from experimental data. F_ij_ can be calculated as in equation (1) using the original data. Theoretically, the position effects of genes i and j are the same, as shown in Equation (5). Assuming that there is no interaction between genes i and j, Equation (6) exists. Therefore, ΔF_i_ (Equation 7) and C_ij_ (Equation 8) can be calculated from Equations (1) to (6). The results are shown in Fig. 2d. If C_*ij*_ > 0, genes i and j have a positive interaction, showing “+.” If C_*ij*_ = 0, genes i and j don’t interact with each other, as shown in white. If C_*ij*_ < 0, genes i and j interact negatively and are displayed as “×.” ΔF_i_ indicates the contribution of a single-gene knockdown to the survival rate of whole-cell libraries.

### Effects of MRP targeting HKs on glycine production

To determine whether the CDCK-SMIA system could function well in MRP of *Vibrio* FA2 via combination interference with the gene expression of CPAs, we selected 35 HK genes as targets for combinational knockdown. Likewise, we used the number of HKs to represent the strains in which the genes corresponding to HKs were knocked down. The cells in the experimental group were divided into two groups based on the eGFP value, Q4 (cell with positive eGFP fluorescence value, defined as FITC-A+) and Q3 (cell with negative eGFP fluorescence value, defined as FITC-A-), with proportions of 95.0% and 4.96%, respectively. Under the same classification conditions, almost all cells in the control group were located in the Q4 quadrant (99.6%). The fluorescence values of cells in the Q4 quadrant were higher than those in the Q3 quadrant. FACS results showed that the percentages of FITC-A+ were 4.85% and 0.24% in the experimental and control groups, respectively, respectively. In summary, these results indicated that glycine production in the cells of the Q4 quadrant might be higher than that of Q3. Thus, the cells in the Q4 quadrant were sorted.

According to the next-generation sequencing (NGS) results, approximately 60,000 reads were obtained per sample. Based on the calculation and statistics between the numbers of reads and dual-sgRNA cassettes, mapping ratios of ∼90.2%, 85.3%, and 80.5% for HK-L1, HK-L2, and HK-S, respectively, were obtained, suggesting that the upstream procedures of CDCK are sufficiently reliable. The frequencies of the different dual-sgRNA cassettes were relatively uniform in the HK-L1 group, whereas some were enriched and some were washed out in the HK-L2 and HK-S groups. After ranking the frequencies of the dual-sgRNA cassettes, the top 3 frequencies of the dual-sgRNA cassette combination in the HK-S group compared to the control were combinational of HK-16 and HK-1 (16-1, 2.0-fold), combinational of HK-32 and HK-2 (32-2, 1.8-fold), and combinational of HK-32 and HK-1 (32-1, 1.7-fold), respectively.

For further validation of the NGS analysis, we independently constructed the strains targeting 16-1, 32-2, and 32-1 (defined as strain 16-1, strain 32-2, strain 32-1), respectively. Thereafter, the cells were cultured under the same conditions (described above) to determine whether the results were consistent with the NGS analysis. The FACS results revealed that the single-cell fluorescence intensities of strains 16-1, 32-2, and 32-1 were 4.2-fold, 4.3-fold, 4.8-fold more than that of the control, respectively. These results were consistent with the NGS data. Furthermore, we extracted the intracellular products of the strains and then measured the amino acid production with an amino acid analyzer. Strain 16-1 was found to produce 353.0 ng/mL of glycine in 20 h under the same culture conditions described above, indicating an increased yield of 3.1% compared to the control strain. Therefore, these results further illustrate that our CDCK-SMIA platform functions well in investigating the effects of MRP**-**targeting HKs on glycine production in *Vibrio* FA2.

### Effects of MRP targeting genes of the glycine pathway on glycine production

To demonstrate the application of the CDCK-SMIA platform in reprogramming the complex metabolic pathway, we selected 24 genes of the glycine pathway as targets, including the genes of the glycine biosynthesis and degradation pathways. The targeted genes were numbered 1−24.

We first constructed a dual-sgRNA library plasmid (defined as G-L1) targeting the glycine pathway using the methods above and then transferred the library plasmid into the *Vibrio* FA2 containing dCas9-pGRS-eGFP plasmid (named G-L2). Cells from the CRISPRi-mediated dual-gene knockdown library of the glycine pathway were screened based on their fluorescence values. Cells with positive fluorescence (defined as G-S) were sorted out through FACS and compared to the control. Thereafter, the plasmids from these cells were extracted and the dual-sgRNA cassette fragments were amplified using PCR. The PCR products were sent for NGS.

We counted the frequencies of different dual-sgRNA cassettes in the surviving cells through NGS data analysis. The frequencies were significantly different among the L1, L2, and S groups. The frequencies of dual-sgRNA cassettes were relatively uniform in the L1 group, whereas some were enriched and some were washed out in the L2 and S groups. This finding may be due to the fact that many cells, in which the core genes of the metabolic pathway were knocked down, did not have the growth advantage over other cells. After careful analysis of the effects of the S group, the dual-sgRNA cassette frequencies of the glycine biosynthesis pathway were found to be markedly higher than those of the degradation pathway. As proof-of-concept, this result is reasonable and consistent with the concept of metabolic engineering ^28, 29^. The dual-sgRNA cassette combination of genes 20 and 13 (20-13, *gcvT*-*ltaE*) had the highest frequency (849 times/10^4^ times). Further, genes 20 and 13 were found to be involved in the glycine degradation pathway, which is consistent with the fundamental theory of metabolic engineering ^28, 29^. Therefore, these results demonstrate that our CDCCK-SMIA platform can be effectively applied in reprogramming complex metabolic pathways.

### Effects of MRP targeting HKs on the antibiotic resistance of *Vibrio* FA2

To test the generality of our CDCK-SMIA plateform for MRP, we changed another HST approach for screening the complex of cell libraries. The CDCK-SMIA platform was used to investigate the effects of MRP on antibiotic resistance of *Vibrio* FA2 when targeting HKs. Instead of screening by FACS based on the fluorescence values described above, we cultivated the strain targeting HKs (HK-L2) in the LB3 medium containing 5 mg/mL ampicillin. 5 mg/mL ampicillin antibiotics caused significant growth inhibition in *Vibrio* FA2. After 16 h of cultivation, the OD_600_ of strain HK-L2 (cultivated with 5 mg/mL ampicillin) decreased by approximately 2-fold compared with that of the control (cultivated without ampicillin). The cells were collected as the growth difference was most remarkable between the control and experimental groups at this time. Independent experiments were conducted three times; the control group was named 1-CK-1, 1-CK-2, and 1-CK-3, respectively; the experimental group was called 1-Ex-1, 1-Ex-2, and 1-Ex-3, respectively. Subsequently, the resulting strains were transferred to fresh LB3 medium as seeds and supplemented with 5 mg/mL ampicillin to strengthen antibiotic screening. The growth difference between the control and experimental group was markedly more significant than that in the previous 28 h. After 33 h of cultivation, the OD_600_ of strain HK-L2 with and without ampicillin was approximately 1.8 and 4.0, respectively. We also harvested the cells for NGS. The experiment was repeated three times; the control group was named 2-CK-1, 2-CK-2, and 2-CK-3; the experimental groups were called 2-Ex-1, 2-Ex-2, and 2-Ex-3, respectively.

SMIA revealed significant differences between the control and experimental groups. According to the NGS data, the top three frequencies of dual-sgRNA cassettes in the experimental group were strains 16-13, 13-29, and 6-33 compared to the control (experiment:control, per 10^4^ times, 272:18, 140:16, and 131:6, respectively). The frequency of dual-sgRNA cassettes of strain 24-30 was 648 in the 2-CK-3 group and 0 in the 2-Ex-3 group. Interestingly, this result was also obtained in another independent experiment, in which the frequency (10^−4^) of strain 24-30 was 323 in the 2-CK-2 group and 0 in the 2-Ex-2 group. The results indicate that ampicillin antibiotic screening enriched some cells (e.g., strain 16-13), and some were washed out (e.g., strain 24-30).

For further validation of the computational analysis, we independently constructed strains targeting HKs of strains 16-13, 24-30, 16, 13, 24, and 30 (the numbers of the HKs were also defined as strains that corresponding HKs were knocked down, e.g. strains 16-13, 24-30, 16, 13, 24, and 30). Thereafter, the strains were cultured in LB3 medium containing the same antibiotic concentrations (described above) to determine whether they complied with the NGS analysis. We found the strains, in which genes HK-16 and HK-13 were simultaneously and individually knocked down, grew faster than the control (wild-type strain expressed dcas9 without sgRNA cultivated in the same condition described above). As expected, these results were consistent with those of the NGS analysis. Interestingly, the *Vibrio* FA2 strain, in which the genes HK-24 and HK-30 were simultaneously knocked down, grew slower than the control. However, the *Vibrio* FA2 strain, in which the genes HK-24 and HK-30 were individually knocked down, grew faster than the control. These results indicate that the metabolic interaction of genes HK-24 and HK-30 may lead to *Vibrio* FA2 decreasing its tolerance to ampicillin antibiotics.

We expressed mRFP in the control strain and eGFP protein in strains 24-30, 16-13, 24, 30, 16, and 13 to further verify these results. Briefly, the strains of experiments and control were co-cultivated under the same conditions described above. Thereafter, the plasmids were extracted from the co-cultured cells at the initial and late logarithmic growth stages and the DNA concentration of the two marker genes were quantified by quantitative PCR (q-PCR). The different DNA concentrations of the marker genes represent the cell numbers of the control and experimental groups. The experimental and control DNA concentrations were almost equal at 0 h. At the late logarithmic growth stages of the co-culture, the DNA concentration of the control was approximately 108-fold higher than that of strain 24-30. In contrast, the DNA concentrations of strains 24 and 30 were was approximately 21-fold and 7.1-fold higher than that of the control, respectively. At the late logarithmic growth stage of the co-culture, the DNA concentration of strain 16-13 was approximately 45-fold higher than that of the control. The DNA concentrations of strains 16 and 13 were also higher than that of the control. The co-cultured bacteria were observed in the late logarithmic stage using a confocal microscope. After simultaneously activating the co-culture cells using FITC, TRITC, and TD channels, the eGFP and mRFP-containing cells were visually observed. The results were consistent with those of q-PCR and the growth curve. Therefore, these results further illustrate that our CDCK-SMIA platform functions well in investigating the effects of MRP on the antibiotic resistance of *Vibrio* FA2.

## Discussion

Synthetic biology is one of the most widely discussed and notable challenges in the 21^st^ century ^30^. As well-known, synthetic biology has been achieving the goal by the rewiring of metabolic networks ^31^, which could be also called as metabolic reprogramming (MRP) via its definition. However, MRP is difficult to study owing to the complexity of the metabolic network, especially from a global metabolic perspective ^32, 33^. Therefore, it is extremely important to operate and remold the metabolic networks.

In this study, we established a MRP platform consisting of CDCK and SMIA that is powerful for operating metabolic networks. We used this platform to harness CPAs for MRP from a global metabolic perspective in *Vibrio* FA2. Our MRP platform can be successfully applied to two aspects: i) interestingly, the effects of MRP targeting HKs and MRP targeting glycine pathway genes on glycine production were investigated; ii) importantly, the effects of MRP targeting HKs on antibiotic resistance of *Vibrio* FA2 were investigated. Moreover, we established a one-stop website for the MRP research, in which users can easily and automatically design related experiments. After wet experiments, the services can be used to conduct data analysis by simply uploading the raw data. In general, it is a smart platform for implementing MRP through the global combinational interference of the CPAs genes.

We successfully conducted this MRP platform via harnessing the cellular perception apparatuses (CPAs), such as HKs, to simulate environmental signal stimulation in *Vibrio* FA2. Naturally existing CPAs are considered to sit on the sensitive nodes of metabolic network, which can trigger natural MRP upon perceiving environmental fluctuations. Traditionally, metabolic engineering has focused on material flow (metabolites) and energy flow (cofactors) ^34^. Therefore, we conducted MRP based on the sensitive nodes, which would be more promising for studying metabolic engineering from the information flow perspective. Using our MRP platform, we identified several genes that are crucial for the ampicillin resistance. Further, we obtained the results of multiple-gene interactions were resulting in changes of metabolic status by only knocking down of several genes via our MRP platform, including the HK genes *sasA_8* and *04288*. Thus, operating these genes of CPAs can be considered as a smart MRP since only manipulating several genes can achieve the goals of the MRP.

Moreover, the genes sitting on sensitive nodes of the metabolic network also include silent genes ^5^ and regulator proteins ^17^, in addition to CPA genes. These apparatuses can be used for the MRP, thereby broadening their applications. Notably, dual-sgRNA libraries were constructed using the gene shuffling method based on PCR, and a dual-sgRNA library targeting n × n genes was obtained by synthesizing only n pairs of primers (n represents the number; Fig. 2a). Thus, our MRP platform is also a relatively simple and cost-effective approach. In addition, the CRISPRi system used in our CDCK approach has been successfully demonstrated to be effective in many hosts, including mammalian cells and bacteria ^23, 27, 35^. Accordingly, our plateform possesses the universal applicability to a wide variety of species.

Controlling metabolic flux is well known to be crucial to achieve the goal of synthetic biology, which has been considered as a “green” technology for producing both bulk and fine chemicals ^36-40^. We attempted to control the metabolic flux to improve glycine production using our method as glycine is an important chemical in the food, pharmaceutical, chemical, agricultural industries ^41^. On one hand, we tested our MRP platform by directly targeting the glycine pathway. According, we successfully demonstrate that knockdown of glycine degradation pathway genes (e.g. *gcvT* and *ltaE*) could increase glycine production. These results were consistent with those of a previous report ^28^, indicating that the MRP platform is reliable. On the other hand, the effects of CPAs on the glycine production by targeting HKs were also investigated. Glycine production was found to be increased by the simultaneous knockdown of the HK genes *cpxA* and *btsS* in *Vibrio* FA2, which was never reported before. The production of biochemicals could also be further improved by interfering with the genes of global regulatory factors (e.g. HKs) in microbial cell factories. Therefore, our MRP is expected to be applied in the biomanufacturing field to obtain promising output.

Metabolic remodeling is also important in microbial physiology; for example, bacterial antibiotic resistance, which is a significant threat to global health, is associated with MRP ^42, 43^. In this study, the effects of HKs on antibiotic resistance were investigated. Several HK genes were found to be related to ampicillin antibiotics in *Vibrio* FA2, such as HK-16 (*cpxA*). This result is consistent with that of a previous study, which was associated with the antibiotics in *E. coli* and *Klebsiella pneumoniae* ^44, 45^. Interestingly, the multiple-gene interactions of the genes HK-24 (*sasA_8*) and HK-30 (*04288*, hypothetical HK protein) could decrease ampicillin antibiotics resistance in *Vibrio* FA2. Moreover, the results were experimentally verified using three methods: q-PCR, growth curve, and confocal microscopy. Based on the results, combinational knockdown can obtain the desired effects rather than independent knockdown of the HK genes *sasA_8* and *04288*. However, the effect of dual-HK genes on antibiotic resistance has never been reported in *Vibrio* strains.The results indicate that the smart MRP platform is also effective at revealing the mechanism of antibiotic resistance, especially the effect of combination genes. Accordingly, our plateform is suggested to be effective for studying the physiology of microorganisms, especially the effects of multiple-gene interactions on antibiotic resistance.

In this study, glycine yield was only slightly increased when investigating the effects of HKs targeting on the glycine production were investigated. The efficiency of high-throughput screening (HTS) methods based on biosensors has been reported to markedly depend on their effects of biosensors ^46-48^. Hence, the slight increase in the glycine production during HKs targeting might be due to the biosensor’s dose-response range of sensing intracellular amino acid concentrations being too narrow. Therefore, the application of our plateform will be further expanded when more effective biosensors become available in the future. On the other hand, Chen et al. put forward the idea that establishing a link between cell growth and products properties allows growth-coupled *in vivo* selection ^49^. Further, Zheng et al. improved amino acid production by coupling amino acid concentration and cell growth ^50^. In this study, growth-coupled screening was successfully applied to investigate the effects of MRP targeting HKs on antibiotic resistance of *Vibrio* FA2. Therefore, the MRP platform would have broader applications when establishing a link between it and growth-coupled *in vivo* selection. Overall, we believe that our approach has high potential and promise.

In summary, we develop a smart MRP platform that include the CDCK and SMIA modes. This platform holds great promise as a versatile smart platform for genome-scale metabolic network reprogramming that is suitable for a variety of fundamental and applied biological research, such as biomanufacturing and drug resistance mechanisms.

## Methods

### Strains, media, and growth conditions

*E. coli* DH5α was used for general cloning and cultivated at 37 °C in a lysogeny broth (LB) medium. *Vibrio alginolyticus* FA2 (*Vibrio* FA2 ^52^) was used for constructing the CRISPRi-mediated dual-gene combination knockdown system and cultivated at 37°C in the LB3 (LB with 3% NaCl, w/w) medium. The *Vibrio* FA2 strain harboring the glycine-riboswitch plasmid was cultivated in the optimized M9 minimal medium ^26^. Chloramphenicol (CmR, 10 μg/mL) or tetracycline (TetR, 5μg/mL) was added to the medium as required.

### Plasmid construction

We used the ClonExpress MultiS One Step Cloning Kit ((Vazyme, Nanjing, China) to construct the plasmids. Golden Gate Assembly was used for designing and constructing the sgRNA library ^51^. The dCas9 and sgRNA chimera genes were synthesized from the previously described vectors ^23^. The dCas9 gene was inserted into the pACYCdute-1 vector containing chloramphenicol-selectable marker and a p15A replication origin, resulting in pdCas9 plasmid. The sgRNA chimera was inserted into the pE1a vector with TetR-selectable marker and ColE1 replication origin, resulting in the pE1a-sgRNA plasmid.

Meanwhile, the two sgRNA chimeras were constructed into pE1a-plasmid, generating the pE1a-2sgRNA plasmid. The pE1a-tool plasmid was constructed by inserting the *ccdB* gene into the N20 position of the pE1a-sgRNA plasmid using the Golden Gate Assembly guideline (Fig. 2a). The mRFP and eGFP genes were synthesized and inserted into the pE1a-sgRNA plasmid to generate the fluorescence marker plasmid of pE1a-mRFP-sgRNA and pE1a-mRFP-eGFP-sgRNA.

### Construction and test of the glycine-on riboswitch

To assess the potential usability of glycine-on riboswitch, different promoters and selective marker gene eGFP were used to construct the plasmids, resulting in the pGRS-pTrc-eGFP (pTrc promoter), pGRS-J23100-eGFP (J23100 promoter), pGRS-Native-eGFP (the promoter of the VC1442 gene in *Vibrio cholerae)*. Then, we deleted the aptamers domain used for binding the glycine in these plasmids, resulting in the control of pGRS-pTrc-Del-eGFP, pGRS-J23100-Del-eGFP and pGRS-Native-Del-eGFP, respectively. All the plasmids were transformed into *Vibrio* FA2 to test whether the glycine-on riboswitch works in *Vibrio* FA2. The strains and their control were cultivated in the optimized M9 medium ^26^ supplemented with various glycine concentrations (0−100 mM). Then, the levels of eGFP fluorescence were determined by FACS.

### Design and construction of the dual-sgRNA library

The primers of the dual-sgRNA library were designed via our one-stop website according to the guideline, which is shown in Fig. 2a. Dual-sgRNA library genes were generated from PCR amplification using the pE1a-2sgRNA plasmid as a template. Then, the PCR products containing the dual-sgRNA library were inserted into the pE1a-tool plasmid using the *BbsI* restriction enzyme and T4 DNA ligase. The complex resulting products were transformed into *E. coli* DH5α competent cells and spread the cells on LB solid medium plate. Then, we washed around 40,000 transformants with LB liquid medium and mixed them as the library 1 strain (L1). The plasmids of the L1 strains were extracted as the dual-sgRNA library plasmid.

### CRISPRi-mediated dual-sgRNA combinational knockdown

The dCas9 protein and pGRS-pTrc-eGFP were constructed into one plasmid. Then, the resulting dCas9-pGRS-eGFP plasmid was transformed into *Vibrio* FA2, defined as dCas9-pGRS-eGFP strain. Next, the dual-sgRNA library plasmid was transformed into the dCas9-pGRS-eGFP strain above. Then, we washed around 40,000 transformants with LB3 medium and mixed them evenly as the initially constructed strain. Then, the arabinose was used to induce the expression of dCas9. Based on their fluorescence values, different cells from the CDCK library were screened using the fluorescence-activated cell sorting (FACS) technology. Compared to the control, cells with higher fluorescence signal fluctuations were sorted out. Next, we extracted the plasmid of these sorted cells as the templates, and the dual-sgRNA fragments were amplified by PCR. Then, the PCR products were sent to NGS ^27^. Hence, the CRISPRi-mediated dual-gene combinational knockdown platform was successfully established.

### Electroporation protocol for DNA transformation

The *Vibrio* FA2 was inoculated into LB3 medium, cultivated at 37 °C, 200 rpm overnight, and then diluted 1:100 into fresh medium. Cells were harvested at OD_600_ nm ∼ 0.5 and pelleted by centrifugation at 4000 rpm for 5 min at 4 °C. The pallet was washed 3 times with pre-cooled buffer (680 mM sucrose, 7 mM K_2_HPO_4_, pH 7.0), and the final cell pellet has resuspended the buffer as a 200-fold concentrate of the initial culture. For transformation, 50 ng plasmid DNA was added into the 100 μL of the competent cell (concentrated cells) in 0.1 mm cuvettes and electroporated using Bio-Rad electroporator at 900 V, 25 µF and 200 Ω and recovered in recovery media (BHI + v2 salt + 680 mM sucrose) for 1h at 37 °C, 200 rpm, then plated on selective medium. The plate was incubated for around 8 h at 37 °C. Brain heart infusion medium (BHI) + v2 salt (1L): 37 g BHI (Becton-Dickinson, Heidelberg, Germany), 204 mM NaCl, 4.2 mM KCl, 23.14 mM MgCl_2_.

### Flow cytometry and analysis and reporter assays

The strains were cultivated in a corresponding medium and contained the appropriate antibiotics at 37 °C, 200 rpm. When the cells were grown to the mid-log phase, the levels of eGFP values were determined by the flow cytometer (BD Biosciences) equipped with a high-throughput sampler. Cells were sampled with a slow flow rate until at least 20,000 cells had been collected. We used the CytExpert software (Beckman Coulter) and Flowjo software (BD) to analyze the FCS data by gating on a polygonal region containing at least 80% cell population in the forward scatter-side scatter plot. All the experiments were measured repeatedly 3 times, and their standard deviation was indicated as the error bar. For mRFP and eGFP assays, the fluorescence was read at excitation/emission of 584/607 nm, and 499/503 nm using the Multifunctional Microplate Reader (Tecan Spark), respectively.

### NGS and data analysis

To analyze the sequencing data, we developed an integrated Python3 software package including sgRNA design and NGS data processing functions (Table S6), because there are no methods available for analyzing our data. The overall workflow of the computational analysis is shown in Fig. 2b. To facilitate the use of this method, we developed a one-stop website, in which the users only need to enter the name of the candidate microorganism, then the well-designed primers are available. The studying candidates of cellular perception apparatus are not only HK but also other genes of two-component systems, silencing genes and regulatory proteins. In short, the convenience and reproducibility of our method were greatly improved.

### Growth curve determination

To assay the effect of ampicillin on *Vibrio* FA2 growth, the strains were grown in the LB3 medium at 37 °C, 200 rpm supplemented with or without 5mg/mL ampicillin (defined as amp^r^+ or amp^r^-group). To assay the *Vibrio* FA2 growth, the OD_600_ of the strains was determined every 5 h using UV–vis spectrophotometer ^52^.

### Microscopy and image analysis

eGFP and mRFP-expressing cells were grown on the LB3 medium until the mid-log phase and observed using the Scanning Confocal Microscope (Nikon Ni-E A1 HD25). The eGFP and mRFP-expressing cells were analyzed by testing each pixel for the presence of FITC, TRITC, and TD fluorescence channels under the same threshold. Images were processed with the image browser and analyzed with the software NIS-Elements Viewer 5.21 64-bit.

### Determination of intercellular glycine concentration

Engineered *Vibrio* FA2 cells and control were harvested under the same OD_600_ nm and disrupted with the extraction buffer (0.1M HCl: 10% (v:v) trichloroacetic acid = 1:2) for 1h. Then the cells were lysed by sonication in the buffer for 15min. The lysate was centrifuged at 10,000 rpm for 5min, and the supernatant was sent to Amino Acid Analyzer (Hitachi L-8900) equipped with column (size: 4.6 mm ID × 60mm, particle size: 3 μm, resin: Hitachi special ion exchange resin).

## Acknowledgments

This work was supported by the grant (2018YFA0903600) from National Key R&D Program of China and by the grants (31870088, 32170105, and 22138007) from National Natural Science Foundation of China.

## Author contributions

Fei Tao and Ping Xu conceived the study. Fei Tao designed the bioinformatical analysis website. Chunlin Tan performed the experiments and analyzed the data. Chunlin Tan wrote the manuscript. Fei Tao and Ping Xu critically reviewed the manuscript and revised it. All authors read and approved the submitted version.

